# High-efficiency CRISPR gene editing in *C. elegans* using Cas9 integrated into the genome

**DOI:** 10.1101/2021.08.03.454883

**Authors:** Matthew L. Schwartz, M. Wayne Davis, Matthew S. Rich, Erik M. Jorgensen

## Abstract

Gene editing in *C. elegans* using plasmid-based CRISPR reagents requires microinjection of many animals to produce a single edit. Germline silencing of plasmid-borne Cas9 is a major cause of inefficient editing. Here, we present a set of *C. elegans* strains that constitutively express Cas9 in the germline from an integrated transgene. These strains markedly improve the success rate for plasmid-based CRISPR edits. For simple GFP insertions, 60 – 100% of injected animals typically produce edited progeny, depending on the target locus. Template-guided editing from an extrachromosomal array is maintained over multiple generations. We have built strains with the Cas9 transgene on multiple chromosomes. Additionally, each Cas9 locus also contains a heatshock-driven Cre recombinase for selectable marker removal and a bright fluorescence marker for easy outcrossing. These integrated Cas9 strains greatly reduce the workload for producing individual genome edits.

**Author Summary:** Germlines have evolved specialized mechanisms to protect themselves from invasions by transposons and viruses, which create barriers to genome editing techniques. For example, transgenes are silenced in the germline of the nematode *C. elegans*, thereby creating a barrier to CRISPR editing by Cas9. To facilitate gene editing, we built a collection of *C. elegans* strains in which Cas9 is never silenced. CRISPR is significantly more efficient in these animals, decreasing the effort researchers need to expend to get edited animals. The strains are available in multiple genetic backgrounds, and contain accessory transgenes to simplify downstream genetics. Together, these strains enable efficient, low-cost genome editing in *C. elegans*.

## Introduction

CRISPR has revolutionized genetics by providing a simple method for targeted mutation, deletion or tagging of genes (Wang et al., 2016). Since its initial demonstration, CRISPR has been rapidly adapted to facilitate gene editing in diverse species. Although the same core set of molecules are used to perform CRISPR editing in different species, methods must be optimized for individual species and applications. Efficient production pipelines are needed to minimize the workload to produce individual edits, and to facilitate systematic production of genome-wide resources.

In the nematode *C. elegans*, the Cas9 endonuclease is introduced into the germline via microinjection of either protein or DNA. In the protein method, Cas9 protein complexed *in vitro* with guide RNA is directly injected into the gonad along with a DNA repair template (Cho et al., 2013; Paix et al., 2015). The injected ribonucleoprotein complex acts immediately in the gonad, producing targeted double-strand breaks within the meiotic oocyte progenitor cells. Co-injected repair templates can be used for homology-directed break repair, yielding edited F_1_ progeny directly from the injected mother. In the DNA method, plasmids expressing Cas9 and the guide RNA are injected into the gonad along with a plasmid DNA repair template (Chen et al., 2013; Dickinson et al., 2013; Friedland et al., 2013). The injected plasmids are assembled into heritable extrachromosomal arrays in the injected gonad. Although cutting is occasionally observed directly in the injected gonad, templated repair events almost always occur in the germline of array-bearing F_1_ animals. Edited progeny are typically recovered among the F_2_ – F_3_ generations.

Direct injection of ribonucleoprotein complexes is fast and efficient – a single injection can yield dozens of independently edited F_1_ progeny (Dokshin et al., 2018; Ghanta and Mello, 2020; Paix et al., 2015). However, scalability is limited by two factors: (1) the reagents are expensive, and (2) edits must be identified by molecular or fluorescence screening of individual progeny because repair template size is limited and selectable markers cannot generally be incorporated. Comparatively, plasmid reagents are inexpensive and can accommodate large repair templates, including selectable markers (Chen et al., 2013; Dickinson et al., 2015, 2013; Norris et al., 2015; Schwartz and Jorgensen, 2016), but array-based edits appear a generation later, editing efficiency is low, and the plasmid-borne transgenes are rapidly silenced. In regard to efficiency, only 5 - 40% of injected P_0_s yield a single edited progeny; most injected animals yield no modified progeny. Although selectable markers allow the rare edits to be identified without molecular screening, the effort required to injecting large numbers of animals limits throughput. Toward developing a pipeline for high-throughput editing in *C. elegans*, we sought to improve the efficiency of plasmid-based CRISPR editing by improving Cas9 expression in worms.

The bane of transgene-driven genome editing in *C. elegans* is silencing of transgenes in the germline. The worm germline detects and limits expression of foreign genetic elements using both sequence-specific and sequence independent mechanisms (Aljohani et al., 2020; Schaner and Kelly, 2006). Silencing of array-borne transgenes is generally incomplete in the F_1_ generation, but strengthens to completely eliminate expression in subsequent generations (Kelly et al., 1997). As the mechanisms of transgene silencing have been elucidated, effective strategies to mitigate silencing have been developed. These include reducing transgene copy number (Frøkjaer-Jensen et al., 2008), increasing array complexity (Kelly et al., 1997), removing pi-RNA complementarity (Zhang et al., 2018), and including Periodic A/T Clusters (PATCs) within introns of the transgene (Aljohani et al., 2020; Frøkjær-Jensen et al., 2016). We have leveraged this knowledge to produce Cas9 transgenes capable of constitutive germline expression.

Here, we present a set of *C. elegans* strains engineered for high-efficiency CRISPR editing using plasmid reagents. The strains harbor integrated, single-copy insertions of germline-licensed Cas9 transgenes. To streamline post-CRISPR strain processing, each Cas9 locus also contains a heatshock-activated Cre recombinase to remove any selectable marker from the edited locus. In addition, each locus is decorated with a fluorescent marker orthogonal to both GFP and mCherry to facilitate removal of the transgene by subsequent outcrossing. These strains greatly improve the efficiency of CRISPR modification using inexpensive plasmid reagents, providing a valuable addition to the *C. elegans* genome engineering toolkit.

## RESULTS

### Single-copy Cas9 transgenes

Plasmid-based CRISPR editing strategies yield poor editing efficiency; typically only ∼20% of successfully injected animals yield edited loci. Therefore to successfully edit with plasmids, many animals must be injected, and broods screened to identify rare events (Dickinson et al., 2015; Norris et al., 2015; Schwartz and Jorgensen, 2016). We found that Cas9 tagged with mCherry was not detectably expressed in the germline in most injected animals, consistent with previous observations (Waaijers et al., 2013). These observations suggest that a lack of germline Cas9 expression is a limiting factor in plasmid-mediated editing. We therefore sought to improve germline Cas9 expression.

We tested whether an integrated single-copy transgene could drive constitutive germline Cas9 expression and generate guided double-strand breaks. We inserted Cas9 transgenes into the ttTi5605 Mos1 transposon insert site on chromosome II using MosSCI (Frøkjaer-Jensen et al., 2008). This locus is known to be licensed for germline expression (Frøkjaer-Jensen et al., 2008). To drive germline-specific expression, Cas9 transgenes were built with the *mex-5* promoter and *tbb-2* 3’UTR. We tested Cas9 efficiency in two strains. In one strain the Cas9 transgene was a cDNA interrupted by a single synthetic intron (derived from Addgene plasmid #46168) (Friedland et al., 2013). In another strain the Cas9 transgene was optimized for germline expression by including 4 large PATC-rich introns from the gene *smu-2* (Fig. 1A)(Aljohani et al., 2020; Frøkjær-Jensen et al., 2016).

**Figure 1.**
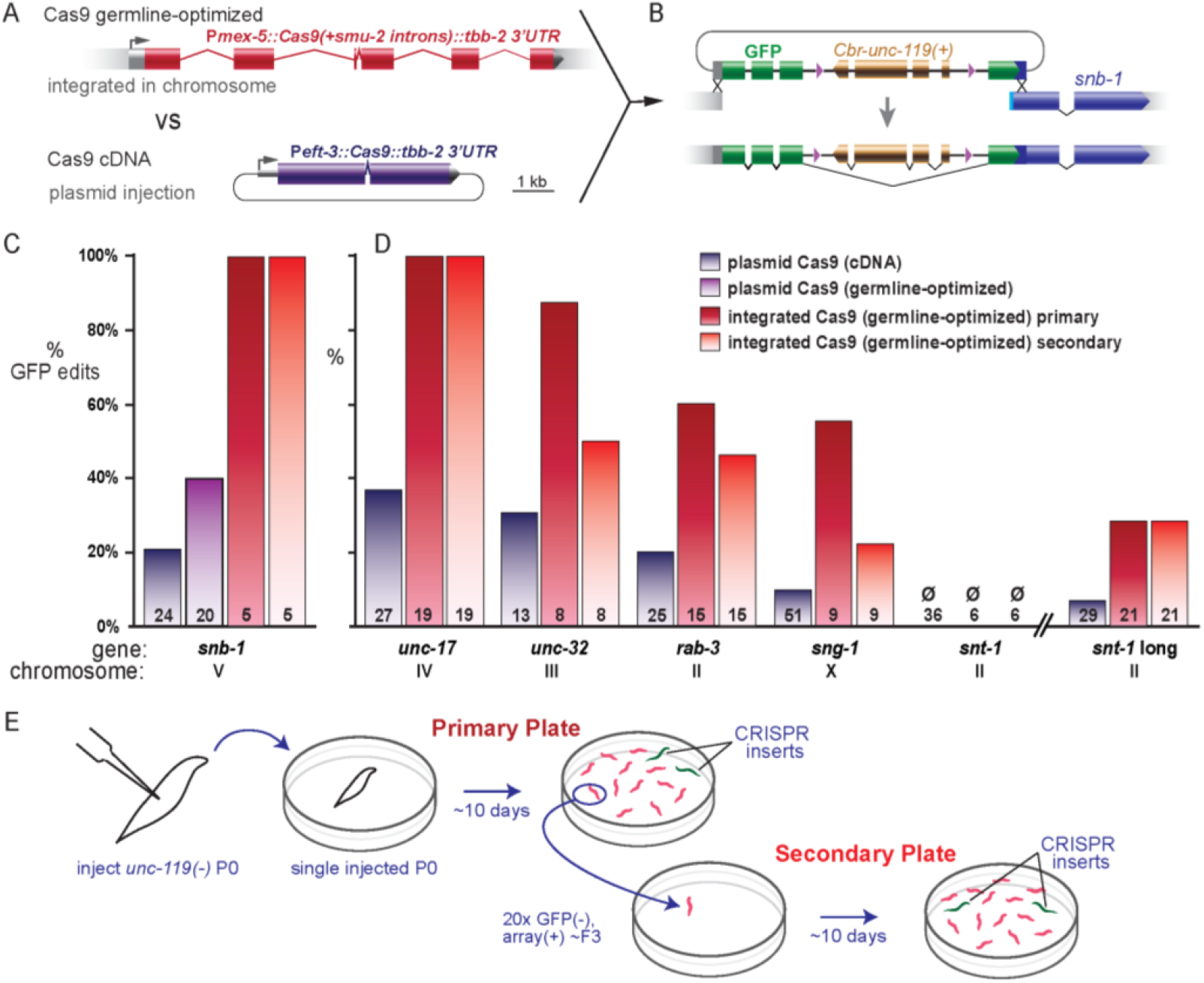
Germline-optimized Cas9 improves editing efficiency. **A**. Gene diagrams representing the germline optimized P*mex-5::Cas9* MosSCI construct and the standard plasmid P*eft-3::Cas9* expression construct. **B**. Diagram of GFP tagging of *snb-1* using the intron-nested *Cbr-unc-119* selectable marker. **C**. Comparison of CRISPR tagging efficiency at *snb-1* using array-borne P*eft-3::Cas9* containing a synthetic intron (dark blue) or germline-optimized introns (light purple) vs integrated, germline optimized P*mex-5::Cas9* (dark red and light red). Insertion frequency is the fraction of successfully injected P_0_s which gave rise to a CRISPR tag among their progeny on the primary or secondary plate (dark red and light red, respectively). Numbers on bars indicate number of successfully injected P_0_s. For germline-optimized Cas9, the tag was always GFP and inserts were confirmed by visual confirmation of GFP expression. For plasmid Cas9, tagging data is reproduced from a previous study with permission (Schwartz and Jorgensen, 2016). For plasmid Cas9 (dark blue and light purple), populations were only incubated until starvation of the primary plate. **D**. GFP insertion efficiency at multiple loci. Editing frequencies were calculated as in (C). **E**. Diagram of multigenerational editing experiment. For integrated Cas9 (dark red and light red), injected P_0_ animals were singled on ‘Primary’ plates. Upon starvation, ‘Primary plates’ were scored for the presence of GFP(+) animals, and 20 array(+), GFP(-) animals were moved to a ‘Secondary plate’. Upon starvation, ‘Secondary plates’ were inspected for GFP(+) animals that arose from new CRISPR events on the secondary plate. Each injected P_0_ was assigned a unique identifier so that editing outcomes for individual lineages in both phases of the experiment could be collated (Table S1).

Each Cas9 expression strain was tested for basal activity by testing for double-strand break formation using the high-efficiency *dpy-10* sgRNA. Animals were injected with a plasmid expressing a *dpy-10* single-guide RNA (sgRNA). We monitored array-positive progeny for dumpy (Dpy) or rolling (Rol) phenotypes consistent with mutation of *dpy-10*. In the strain bearing the germline-optimized Cas9 construct (EG9615), ∼25% of array-bearing F_1_ animals (18/69) exhibited a Dpy or Rol phenotype. To determine if these were heritable mutations, we singled F_1_ animals. Roughly half of these (8/15 singled F_1_s) were caused by germline mutations, that is, the singled animals produced either 100% Dpy or one quarter Dpy progeny, consistent with germline mutation. By contrast, in strains expressing the Cas9 cDNA construct, we failed to detect *dpy-10* mutant animals among similar numbers of array positive progeny; this strain was not characterized further. Together, these data indicate that a single-copy transgene can express sufficient Cas9 to introduce cuts in the genome, but it must be licensed for germline expression.

As a more demanding test of Cas9 activity, we knocked a GFP tag into the N-terminus of SNB-1 (Fig. 1B). GFP insertions were selected by co-insertion of an *unc-119(+)* transgene nested in an intron of GFP (Schwartz and Jorgensen, 2016). Editing efficiency was compared to plasmid injection of the germline-optimized Cas9 transgene, as well as injecting plasmid expressed Cas9 cDNA (Schwartz and Jorgensen, 2016). To normalize for variable injection quality, genome edits were only quantified from injected animals that generated heritable extrachromosomal arrays and were therefore verified as ‘successfully injected’. Edits were observed in 100% of successfully injected P_0_s using the integrated germline-optimized Cas9 transgene (Fig. 1C), which is a profound improvement over our results using plasmid-injected cDNA (20% P_0_s). Injecting the germline-optimized Cas9 transgene as a plasmid improved injections marginally (40% P_0_s) and was silenced thereafter; this construct was not characterized further. The integrated transgene was then tested on a variety of genes to determine if it consistently outperformed plasmid injection of the cDNA.

We injected targeting plasmids specifying GFP insertion at five other synaptic genes previously used to assay insertion frequency using array-borne Cas9 (Schwartz and Jorgensen, 2016). For each locus, the targeting reagents comprise a repair template plasmid and a guide RNA expression plasmid. All genes were initially targeted with “short-arm” repair templates (≤ 57 bp) assembled by SapTrap (Schwartz and Jorgensen, 2016). The *snt-1* gene was additionally targeted by a second “long-arm” construct (150 and 450 bp). Across all loci, GFP insertion per successfully injected P0 was 2.5- to 7-fold more frequent in the integrated Cas9 strain than in strains expressing Cas9 from extrachromosomal arrays (Fig. 1D). The short-arm *snt-1* targeting strategy uses a guide RNA predicted to exhibit low activity due to a “C” base immediately preceding the PAM site (CNGG). The enhanced activity from integrated Cas9 was unable to overcome this guide RNA deficiency; whereas successful integration was observed for an upstream cut site requiring a long-arm template. For all other short-arm constructs, between 50% and 100% of injected P0’s yielded tagged inserts. Thus, the integrated Cas9 transgene facilitates more efficient CRISPR editing than array-borne Cas9.

We next explored whether the integrated Cas9 transgene could support editing over multiple generations. In general, array-based genome editing in *C. elegans* is limited to a few generations after the initial injection, even though the editing arrays can transmit indefinitely. This is likely due to silencing of the Cas9 transgenes on the extrachromosomal arrays. The integrated and germline-licensed Cas9 is not silenced; all of the GFP targeting constructs in Figure 1 were injected in an integrated Cas9 strain more than ten generations after the initial integration of Cas9. Although the integration provides continuous Cas9 expression, it was not known if array silencing mechanisms would limit guide RNA expression or repair template availability after the first few generations. We therefore monitored our GFP-tagging injections for new editing events beyond the F_2_ generation, when germline array silencing is usually established (Kelly et al., 1997).

To detect new CRISPR events in later generations, we propagated unedited, array-bearing worms past the F_3_ generation (that is, beyond starvation of the primary plate). Successful edits are easily scored by bright GFP expression in the synapse-rich nerve ring, which allowed us to reliably pick unedited individuals using a fluorescence dissecting microscope. Upon starvation of each plate founded by a single injected P_0_ animal, we singled 20 animals that carried the array (*unc-119+*) but lacked an edited target locus (GFP-). When these ‘secondary’ plates starved (approximately 3 generations), the plates were inspected for GFP(+) worms (Fig. 1E). For all except the defective short-arm *snt-1* target, new GFP insertions occurred on the secondary plates, albeit usually at a slightly lower rate than on the primary plates (Fig. 1D). In most cases, lineages that generated inserts on the secondary plate had produced CRISPR insertions on primary plate (Table S1). Interestingly, four lineages that failed to generate CRISPR inserts on the primary plate nevertheless yielded GFP inserts on the secondary plate. Thus, in the integrated Cas9 strain, *de novo* editing events routinely occur in generations after array silencing. Importantly, polymerase III-mediated guide RNA expression and repair template accessibility are less sensitive to germline silencing mechanisms than polymerase II-mediated Cas9 expression.

### Optimizing Cas9 expression strains

Since integrated Cas9 transgenes delivered significantly higher success rates in GFP tagging experiments, we developed optimized strains to streamline the editing pipeline for the *C. elegans* community. These Cas9 strains include:

1. A fully outcrossed and uniform genetic background for *unc-119* rescue.
2. Cas9 integrations on different chromosomes.
3. A linked, inducible Cre recombinase to excise the selection marker in the edited locus.
4. A fluorescent marker in the Cas9 locus to facilitate out-crossing.
5. At least one Cas9 integration tightly linked to *unc-119*, so that the selectable marker is outcrossed simultaneously with the Cas9 transgene.

To generate a uniform genetic background, we thawed the standard N2 strain and generated a *de novo unc-119* mutation. We injected plasmids encoding Cas9 and a guide RNA targeting the region near the *ed3* allele of *unc-119*, and relied on error-prone repair to generate a new loss-of-function allele. We isolated a single *unc-119(-)* strain and designated the new allele *unc-119(ox819)*. Direct sequencing of *unc-119* revealed *ox819* is an 11 bp deletion flanking the targeted cut site (Fig. S1). *ox819* causes a frameshift after V110 (UNC-119a) and then appends 29 out-of-frame amino acids before a stop codon. *unc-119(ox819)* animals are phenotypically identical to *unc-119(ed3)* animals and can be rescued by expression of the smaller *unc-119* gene from the related nematode *C. briggsae* (*Cbr-unc-119*). The strain carrying *unc-119(ox819)* represents a relatively clean genetic background for gene editing since it was generated by targeted rather than random mutagenesis to limit secondary mutations, and the strain was immediately frozen to minimize the emergence of suppressor mutations during propagation of the strain in laboratory stocks.

To insert Cas9 expression cassettes at a variety of loci in the genome, we randomly integrated the germline optimized Cas9 expression cassette into the *unc-119(ox819)* background using a miniMos transposon (Fig. 2). The transposon carried a Cas9 expression cassette, P*mex-5::Cas9* (+*smu-2* introns)::*tbb-2* 3’UTR, and a lox-flanked *unc-119(+)* locus for selection. We isolated 14 inserts and mapped the miniMos insertion site by inverse PCR (Table S2). We first assessed germline Cas9 activity by confirming the ability to cut the *dpy-10* locus. We injected the plasmid expressing the *dpy-10* sgRNA and assayed for the presence of Dpy or Rol animals. Second, we assayed for efficient CRISPR-mediated GFP insertion into the *unc-32* gene (Table S2). This screen identified four strains in which (1) Cas9 activity was high enough for robust targeted GFP insertion (> 50% of injected P0’s gave rise to *unc-32::GFP* progeny), and (2) the Cas9 expression cassettes were distributed broadly across autosomes. In three of the four strains, the Cas9 expression cassettes were inserted in unnamed genes without exacerbating phenotypes. Remarkably, in the fourth strain, the transposon (*oxTi1083*) landed in the *unc-119(ox819)* locus on chromosome III, satisfying the goal of a Cas9 integrant tightly linked to *unc-119*. The screen identified at least one strain (inserted into *Y69H2*.*3* on chromosome V) capable of editing *dpy-10* but not capable of supporting homology-dependent GFP tagging of *unc-32* (Table S2). Together, our screen results are consistent with previous observations that the local genomic environment can influence the overall level of germline expression (Frøkjær-Jensen et al., 2016), and that different levels of germline Cas9 expression can lead to varying editing efficiency.

**Figure 2.**
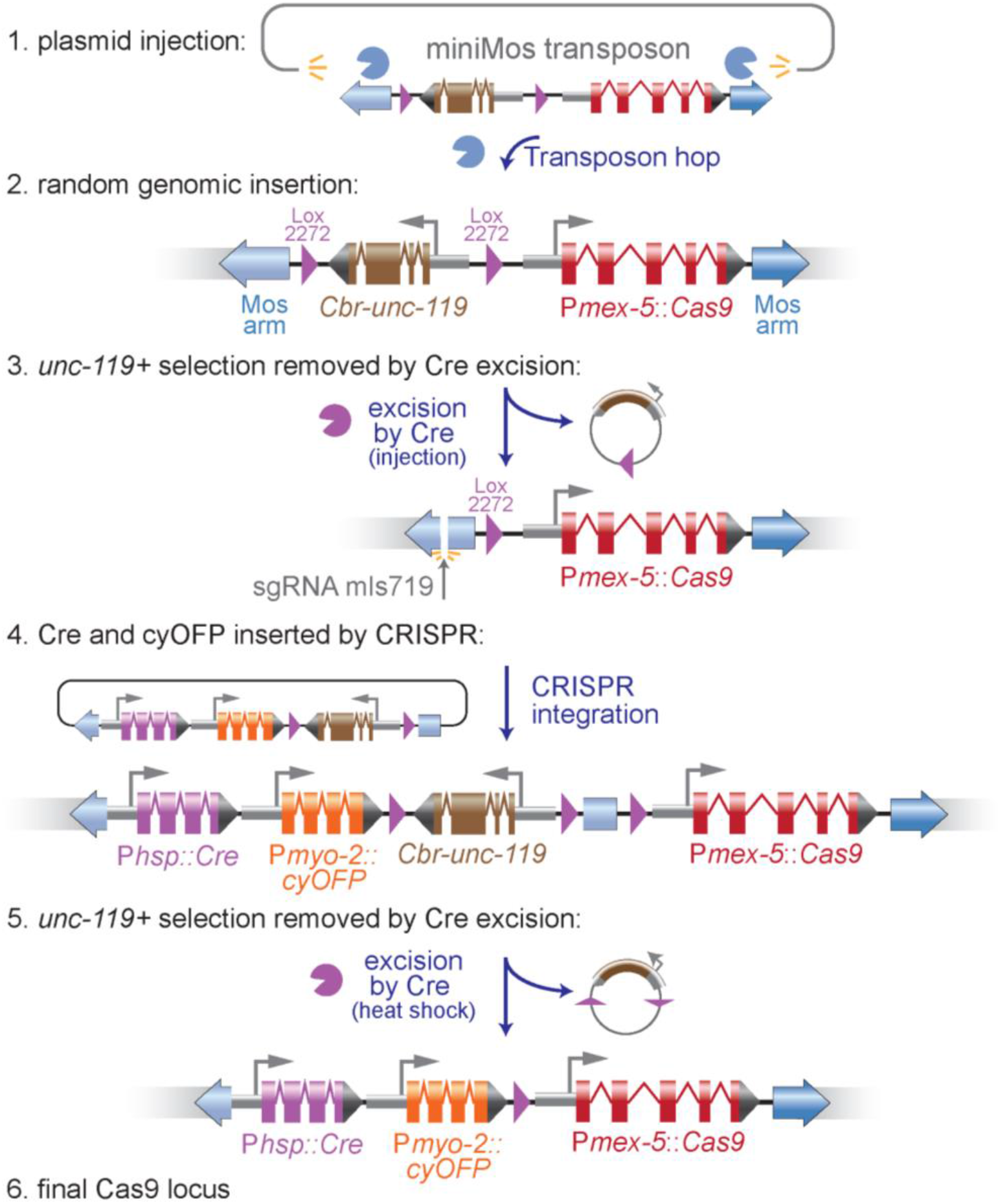
Building marked Cas9-Cre loci. (1) Cas9 transgenes were generated by injecting a miniMos transposon containing a germline-optimized Cas9 and a floxed *Cbr-unc-119* selectable marker into *unc-119(ox819)* mutants. (2) MiniMos elements inserted into the genome randomly and injection sites were mapped by inverse PCR and confirmed by directed PCR. (3) The *Cbr-unc-119* cassette was removed by injecting a plasmid expressing Cre. (4) The loci were then modified by CRISPR to include a Cre transgene and an Orange Fluorescent Protein (shown), or just a Cre transgene (not shown). The cut site was introduced by injecting a plasmid expressing a guide RNA for a site in the 5’ arm of the Mos1 transposon (sgRNA mls719); Cas9 was expressed by the transgene. The insert included P*hsp-16*.*41::Cre* + P*myo-2::cyOFP* (or *Phsp-16*.*41::Cre* alone) and a floxed *Cbr-unc-119* selectable marker. (5) Strains were heat shocked to activate Cre and remove the *Cbr-unc-119* marker, yielding the final strains (6) in an *unc-119(ox819)* background.

We next incorporated a Cre locus and fluorescent marker to each locus to simplify genetic manipulations. To modify any of the miniMos insertions, we built a tagging construct that targets insertions into the 5’ arm of the miniMos element (Fig. 2, step 4). The targeting vector contains a lox2272-flanked *Cbr-unc-119* cassette to select for the modified miniMos insertion. After insertion, heat-shock activation of Cre removes the *unc-119(+)* transgene.

To each of the Cas9 loci we added a heatshock-driven Cre transgene (P*hsp16*.*41*::Cre::*tbb-2* 3’UTR). For tagging genes, we use a selection strategy in which a floxed *Cbr-unc-119* is nested in an intron of the inserted tag, thereby rescuing the *unc-119(ox819)* mutation at the genomic locus. The *Cbr-unc-119* gene is excised from the edit, leaving only a loxP site within an intron in GFP. Integrated heat-shock Cre eliminates the need for a secondary injection of the Cre-expressing plasmid to excise the selectable marker and drives more efficient marker excision (Dickinson et al., 2015).

To simplify outcrossing of the Cas9 locus, we generated additional strains that incorporated a fluorescent marker. In these cases the CRISPR template contained transgenes for both Cre and an orange fluorescent protein (P*myo-2*::*2xNLS-cyOFP*::*let-858* 3’UTR). cyOFP (cyan-excitable Orange Fluorescent Protein) is a long-Stokes-shift fluorescent protein that is spectrally separable from common green and red fluorophores. The nuclear localized, *Pmyo-2-*driven transgene produces a bright fluorescence pattern that is limited to the pharynx (Fig. S2); the marker is brightest in late larvae (L3 and L4) but dimmer in adults, providing a marker that is easy to track through crosses but that minimally obscures weak fluorescence signals from tagged target genes.

We quantified the activity of both Cre and Cas9 in each of the final strains. We assayed Cre activity by monitoring the efficiency of excision of the *Cbr-unc-119+* marker from the locus (Fig. 2, step 4). Heatshock-driven Cre acts efficiently from integration sites in the self-excising cassette (SEC) platform (Dickinson et al., 2015). We found that heatshock-driven Cre excised *Cbr-unc-119+* in ∼50% of the progeny of heat-shocked animals (Fig. 3A). We observed a low amount of Cre activity from the heat-shock transgene even in the absence of a *bona fide* heat-shock treatment. This ‘leaky’ Cre expression leads to marker excision events when animals are reared at 20 – 25°C but is mostly eliminated by rearing animals at 15°C.

**Figure 3.**
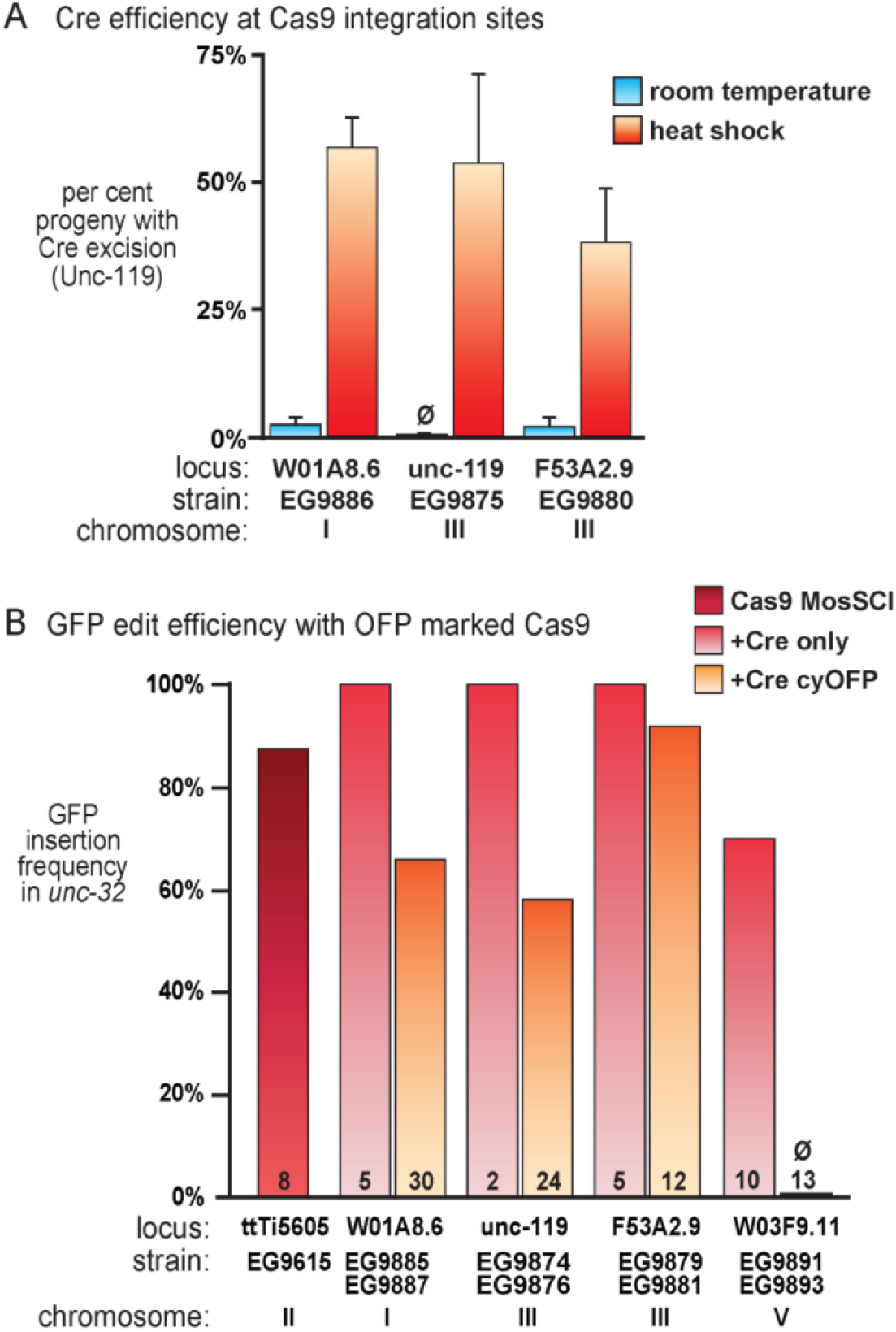
Cre and Cas9 activity from integrated Cas9 loci. **(A)** *unc-119(+)* marker excision using integrated heat-shock Cre transgene. Cohorts of ∼20 homozygous Unc-119(+) young larvae (L1 – L2) were either incubated at room temperature or at 32°C for 2 hours (“heat shock”). After reaching adulthood, several hundred eggs (n = 223 – 1025) were collected from each cohort, and each hatched animal was scored for the presence of Unc-119(-) phenotype. Bars represent standard deviation for 3 independent experiments. (**B)** Insertion of GFP into the *unc-32* gene using CRISPR. The *unc-32* guide RNA was expressed from an injected plasmid; Cas9 was expressed from a transgene integrated at the indicated locus. A *Cbr-unc-119(+)* transgene is contained within an intron of the GFP tag. The loci either contained the Cre transgene alone (light red bars, top strain name) or the Cre and the Orange Fluorescent Protein transgenes (orange bars, bottom strain name). The first strain (EG9615) contains just the Cas9 transgene at the MosSCI insertion site (ttTi5605). Configurations of the transgene loci in these strains is illustrated in Figure S4. Insertion frequency is the fraction of successfully injected P_0_s which gave rise to a CRISPR tag insertion among their progeny. Numbers on bars indicate injected P_0_s, chromosome indicates the location of the insertion containing the Cas9-Cre locus.

After excision of the *Cbr-unc-119* marker from the Cas9-Cre locus, we quantified Cas9 activity in each strain by measuring the insertion frequency of *unc-32::gfp* (Fig. 3B). The Cas9 insertions on chromosomes I and III maintained moderate-to-high levels of Cas9 activity. Unexpectedly, the Cas9 insertion on chromosome V (*W03F9*.*11*) was silenced by the addition of the cyOFP transgene. Multiple independent insertions of P*myo-2*::*2xNLS-cyOFP*::*let-858 3’UTR* into the chromosome V site silenced Cas9 activity. Additionally, insertion of P*myo-2*::*2xNLS-cyOFP*::*let-858 3’UTR* up to 0.3 map units away from the Cas9 transgene resulted in silencing of the Cas9 transgene (not shown). Thus, we were only able to produce fluorescently marked alleles for the chromosome I and chromosome III insertions. However, Cas9 expression remains robust in a simple version with just the Cre transgene inserted in the chromosome V Cas9-Cre locus.

Finally, to facilitate use with selectable markers other than *Cbr-unc-119*, we outcrossed the fluorescently marked chromosome I (W01A8.6) and chromosome III (F53A2.9) Cas9-Cre transgenes to a wild-type background. Because one of the Chr III inserts disrupts *unc-119* directly, this allele could not be segregated from an *unc-119* mutant background. Together, we have produced a suite of Cas9 and Cre expressing strains capable of supporting efficient plasmid and array-based CRISPR editing in *C. elegans*.

## Discussion

In summary, we have generated specialized strains for efficient genome editing by CRISPR. These strains express the Cas9 enzyme in the germline constitutively, and can insert tags in a variety of genes at a success rate of 60%-100% of injected worms.

There are four important features to these strains that permit high expression of Cas9. First, to avoid silencing of the construct, Cas9 is expressed from an integrated transgene rather than from an injected plasmid. In *C. elegans* injected DNA forms extrachromosomal arrays, which are usually silenced in the germline (Kelly et al., 1997; Mello et al., 1991; Stinchcomb et al., 1985). Second, we inserted introns containing periodic A/T clusters (PATCs) into the Cas9 transgene. These clusters are simply a few adjacent A/T basepairs present with a 10 bp periodicity on a single face of the DNA molecule (Fire et al., 2006). Inclusion of introns containing PATCs facilitates expression of genes in the *C. elegans* germline (Aljohani et al., 2020; Frøkjær-Jensen et al., 2016). This specialization may protect the germline from invasion by transposons or viruses. Because foreign DNA is unlikely to contain these specialized introns, they will not be expressed in the germline. Third, because not all regions of chromosomes are permissive for germline expression, we tested 15 chromosomal integration sites, and 5 of these insertions expressed sufficient levels of Cas9 for efficient gene tagging with GFP. Fourth, we labelled 4 of these sites with a fluorescent marker and retested tagging efficiency; 3 of the 4 sites retained high Cas9 expression with the presence of the fluorescent marker.

The immediate advantage of the Cas9 expression strains is that they greatly reduce the injection burden for producing CRISPR edits with plasmid reagents. Previously, plasmid-based protocols required ∼five times more injection effort than direct injection of the Cas9 protein. The significant improvement in efficiency afforded by the Cas9 expressing strains brings the injection burden for plasmid reagents on par with protein injections (Ghanta and Mello, 2020). Cas9 ribonucleoprotein complex injections are likely to remain the preferred method for short edits including point mutagenesis. For tagging and gene-deletions that require long repair templates, the enhanced efficiency of Cas9 expressing worms coupled with selection within the tag will make plasmid-based editing an important alternative.

In previous DNA protocols Cas9 was expressed from a plasmid; edits largely occurred in the F1 germline only, apparently because the transgene in the extrachromosomal array is silenced thereafter. When Cas9 is expressed from an integrated transgene, editing is not silenced in subsequent generations, and unedited strains that carry the array will continue to yield GFP insertions in later generations. Continued editing capability suggests that the guide RNA plasmid expressed by a U6 RNA polymerase III promoter is still active on the extrachromosomal array, and the DNA repair template with the GFP tag is still a substrate for double-strand break repair of the cut site.

Nevertheless, there is variability in editing efficiency among our standard set of test loci for gene tagging, but this variability is unlikely due to germline silencing. The three genes that were edited most efficiently on both primary and secondary culture plates are essential genes, whereas the three genes that were edited less efficiently are non-essential. One potential explanation for this observation is that the target sites are being cut repeatedly until either a templated or untemplated mutagenic repair occurs. For essential genes, the untemplated repairs result in lethal alleles that are lost from the population. For non-essential genes, untemplated repairs create viable mutations that lack the targeted cut site. Indeed, limited sequencing of ‘dead-end’ strains revealed mutated guide RNA binding sites (not shown).

We have found two primary modes in which our non-Unc non-array screening fails to identify correct genome edits using our strains: (1) ‘dark arrays’ and (2) ‘dark inserts’. Both these modes mimic *bona fide* inserts, as they have *Cbr-unc-119(+)* rescue but not fluorescent array markers. A dark array is an array that lacks detectable fluorescent markers, either because the markers were never incorporated or because the markers were excised or silenced during propagation. These animals are selected because they exhibit Unc-119 rescue but lack array fluorescence. Lines harboring dark arrays generally become apparent in the F1 of selected animals; whereas genomic inserts are transmitted to 75% or 100% of the progeny through chromosomal segregation, array segregation is generally much less efficient. Arrays lacking one or more injected plasmids and thereby mimic genomic edits have been observed in other forms of array-based genome editing (‘false positives’, wormbuilder.org). Dark inserts, on the other hand, are insertions in the genome in which GFP is not expressed. These are due to mutated or partial edits, or off-target insertions of the *Cbr-unc-119(+)* cassette into the genome. Through whole genome sequencing we have characterized a small number of dark, off-target inserts, and they appear to be inserted non-specifically in the genome, likely into off-target Cas9 cut sites or other double-strand breaks (data not shown). Dark inserts lack array markers and are rescued for *unc-119*, but do not express GFP and the genotype cannot be confirmed by PCR. Of 206 lines fully analyzed in this study, we identified 3 plates containing animals lacking array markers but also lacking a GFP insert. Thus, the observed rate of these failure modes is below 5% of injected P0’s. Neither dark arrays nor dark inserts are specific to integrated Cas9. Both are observed at similar frequencies when Cas9 is expressed from an injected plasmid. Importantly, both of these failures modes can be easily identified and excluded from further processing.

To build a versatile genetic toolkit for CRISPR modifications, we inserted Cas9 transgenes at various sites in the genome. We tested 15 integration sites, 5 of these insertions expressed sufficient levels of Cas9 for efficient gene tagging with GFP and were on diverse locations in the genome (Fig. S3). Four of these loci were modified to include a Cre transgene to remove the selection marker from the edited gene, and 3 modified to include a fluorescent marker so that the Cas9 transgene could be easily outcrossed to generate the final edited strain (Fig. S4). We include a protocol to illustrate the steps for successful gene editing and outcrossing (Fig. S5).

## Methods

### Reagents

All chemicals were purchased from Sigma-Aldrich (St. Louis). All enzymes were purchased from New England Biolabs (Beverly, MA). All synthetic DNAs were purchased from Integrated DNA Technologies.

### C. elegans strains

*C. elegans* strains were cultured using standard methods (Brenner, 1974) on nematode growth media (NGM) feeding on OP50 or HB101 bacteria. Animals were maintained at 15°C, 18°C, and 20°C. A list of worm strains created in this study can be found in Table S3.

### Molecular biology and cloning

A list of plasmids used in this study can be found in Table S4. All novel plasmids were generated using standard molecular biology techniques. Annotated sequences for all plasmids used in this work are included in Supplemental File 1.

#### General Injection Procedures

All injections were conducted into young adult (<24 hours) hermaphrodites reared at 15°C on either OP50 (wild type worm strains) or HB101 (*unc-119* mutant worm strains). For plasmid injections, plasmids were purified using the *Purelink* kit (Thermofisher). Unless noted otherwise, the final total concentration of all injected plasmids was 100 ng/µl. Arrays were marked by inclusion of either 1xRed co-injection markers [2 ng/µl pCFJ90 (P*myo-2::mCherry)*, 4 ng/µl pGH8 (P*rab-3::mCherry*) + 4 ng/µl pCFJ104 (P*myo-3::mCherry)*] or 1xGreen co-injection markers[2 ng/µl pCFJ91(P*myo-2::gfp*) + 8 ng/µl pCFJ421 (P*eft-3::gfp::h2b*)]. For CRISPR injections with a single plasmid containing both the repair template and guide RNA expression cassette, the plasmid was included at 65 ng/µl. For CRISPR injections with the repair template and guide RNA expression cassette on different plasmids, the repair template was included at 60 ng/µl and the guide RNA expression plasmid was included at 30 ng/µl. When included, Cas9 expression plasmids were included at 30 ng/µl (cDNA) or 2 ng/µl (with smu-2 introns); higher levels of the germline-licensed Cas9 transgene were toxic. If needed to bring injection mix concentrations to 100 ng/µl, pBluescript(sk+) was used as “stuffer” DNA. After injection, P_0_s were either singled to fresh OP50 plates (when quantifying event frequency per injected P_0_) or pooled in groups of 3 – 5 (for construction operations) and incubated at 25°C for 3 – 10 days prior to screening.

#### Transposon-mediated Cas9 insertion

All MosSci injections were conducted with young adult EG6249 worms. MosSci injection mixes contained 25 – 40 ng/µl ttTi5605-targeting vector, 50 ng/µl pCFJ601 (P*eft-3::Mosase*), 10 ng/µl 1xRed markers, and 10 ng/µl pMA122 (P*hsp-16*.*41::peel-1*) (95 - 110 ng/µl total). MiniMos injections were conducted with young adult EG9814 *[unc-119(ox819) III]* worms. MiniMos injection mixes contained 50 ng/µl pCFJ601 (P*eft-3::Mosase*), 40 ng/µl pMLS714 (miniMos targeting vector), and 10 ng/µl 1xRed markers. Initial MosSci inserts (oxSi---- and oxSi1091) were built using the standard pCF150 targeting vector. To allow removal of the *Cbr-unc-119* removal after transgene insertion, we built general targeting vectors for MosSci at the ttTi5605 locus (pMLS640) and for MiniMos (pMLS713) containing a *lox2272-*flanked *Cbr-unc-119* cassette. In all cases, genomic inserts were identified by screening for Unc-119+ animals that lacked extrachromosomal array markers. The floxed *Cbr-unc-119* was removed from strains by injecting an array containing 50 ng/µl pDD104 (P*eft-3::Cre)* and selecting for *unc* F_2_ progeny.

#### Cas9 locus modification

Cas9+, *unc 119-*animals were injected with 90 ng/µl pMLS719 (miniMos-targeting sgRNA), 10 ng/µl either pMLS716 (P*hsp-16*.*41::Cre + lox2272* flanked *Cbr-unc-119*) or 2 ng/µl pMLS791 (P*hsp-16*.*41::Cre +* P*myo-2::2x-nls-cyOFP + lox2272* flanked *Cbr-unc-119*) + 10 ng/µl 1xRed marker. Cas9 activity was provided by the integrated Cas9 transgene. Injected P_0_ were incubated at 25°C for 10 days. Inserts were selected by *unc-119* rescue and the absence of fluorescence array markers.To remove the *Cbr-unc-119* marker, larval animals were heat shocked at 32°C for 2 hours. *Unc-119-*animals were selected by phenotype from the F_1_ progeny of heat-shocked animals.

#### Measuring Cas9 activity with dpy-10 sgRNA

For initially screening MosSci Cas9 strains, worms were injected with a low concentration of *dpy-10* sgRNA plasmid to improve our dynamic range: 10 ng/µl pMLS597 (P*U6::sgRNA*(*dpy-10*)), 10 ng/µl 1xRed markers, and 80 ng/µl stuffer DNA. For screening miniMos-based Cas9 strains, worms were injected with 90 ng/µl pMLS597 (P*U6::sgRNA*(*dpy-10*)) and 10 ng/µl 1xRed markers. The higher concentration of pMLS597 resulted in a higher rate of germline mutagenesis of *dpy-10* among array(+) animals (not shown). Injected P0 animals were pooled on fresh OP50 plates and incubated at 25°C for 2 – 3 days. Array(+) F_1_ animals were selected by fluorescence from the co-injection markers. Selected worms were later scored for Dpy and Rol phenotypes under white light.

#### Measuring tag (GFP) insertion frequencies

For quantifying insertion frequency at multiple target loci in EG9747, worms were injected with 65 ng/µl targeting vector, 10 ng/µl 1xRed markers, and 25ng/µl suffer DNA. For quantifying insertion frequency in miniMos insertion strains, worms were injected with 90 ng/µl targeting vector (pMLS338) and 10 ng/µl 1xRed markers. We used only 65 ng/µl targeting vector when quantifying insertion frequency in EG9747 so that the targeting vector concentration in these injections matched the targeting vector concentration used for quantifying insertion frequency at these sites using plasmid-borne Cas9 (Schwartz and Jorgensen, 2016; data reproduced with permission). After injection, P0 animals were singled to OP50 plates and incubated at either 25°C or room temperature (20°C) until starvation. Starved plates were inspected for array+ and GFP+ animals. Insertion frequency was calculated as (number of plates with GFP + worms) / (number of successful injections), where a ‘successfully injection’ is an injection resulting in array+ worms in the F_2_ generation.

#### Quantifying floxed marker excision rates

For quantifying Cre activity, we used *unc-32::GFP(Cbr-unc-119+)* alleles created in either the EG9886, EG9875, or EG9880 background. Cohorts of ∼20 homozygous young larvae (L1 – L2) of each strain were either incubated at room temperature or at 32°C for 2 hours (“heat shock”). After adulthood, eggs were collected from each cohort, and each hatched animal was scored for the presence of Unc-119(-) phenotype.

## Acknowledgments

We thank Christian Frøkjaer-Jensen for suggestions on licensing germline transgenes. We thank the Caenorhabditis Genome Center for maintaining and distributing strains. Siyu Chen provided pSYC33, Brian Mueller and Adam Hefel vetted the manuscript, and members of the Jorgensen lab provided vigorous discussions. EJ is an Investigator of the Howard Hughes Medical Institute. This work was funded by NIH grant R01GM095817. MSR was supported by NIH grant F32GM133139.

## Supplemental Figures

**Supplemental Figure 1.**
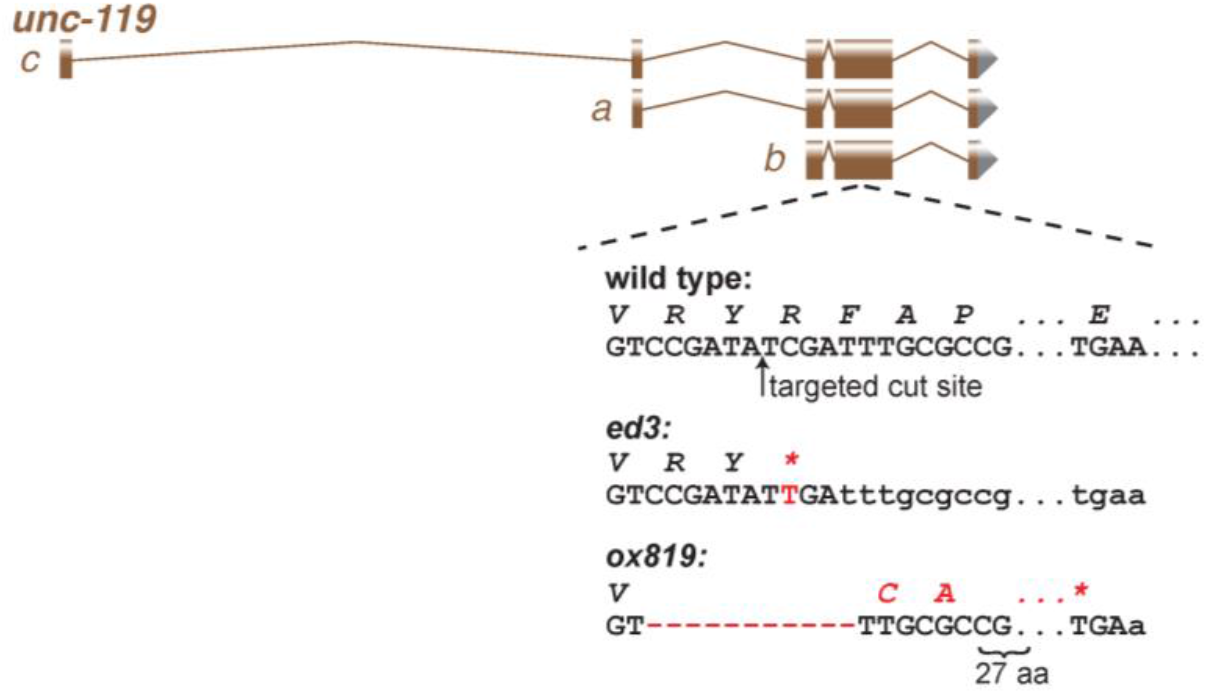
Generating *unc-119(ox819)*. The classic *unc-119(ed3)* allele was generated using EMS, and is likely to have other mutations in its background. Because CRISPR edits should be generated in a wild-type background that has not been exposed to a chemical mutagen, we generated a new *unc-119* mutation in a wild-type background. N2 worms were injected with a plasmid encoding Cas9 and a guide RNA targeting the same region of *unc-119* as *ed3*. Sequencing of *ox819* revealed an 11 base-pair deletion starting at V110 (UNC-119a) straddling the targeted cut site which leads to a frameshift and stop after 27aa.

**Supplemental Figure 2.**
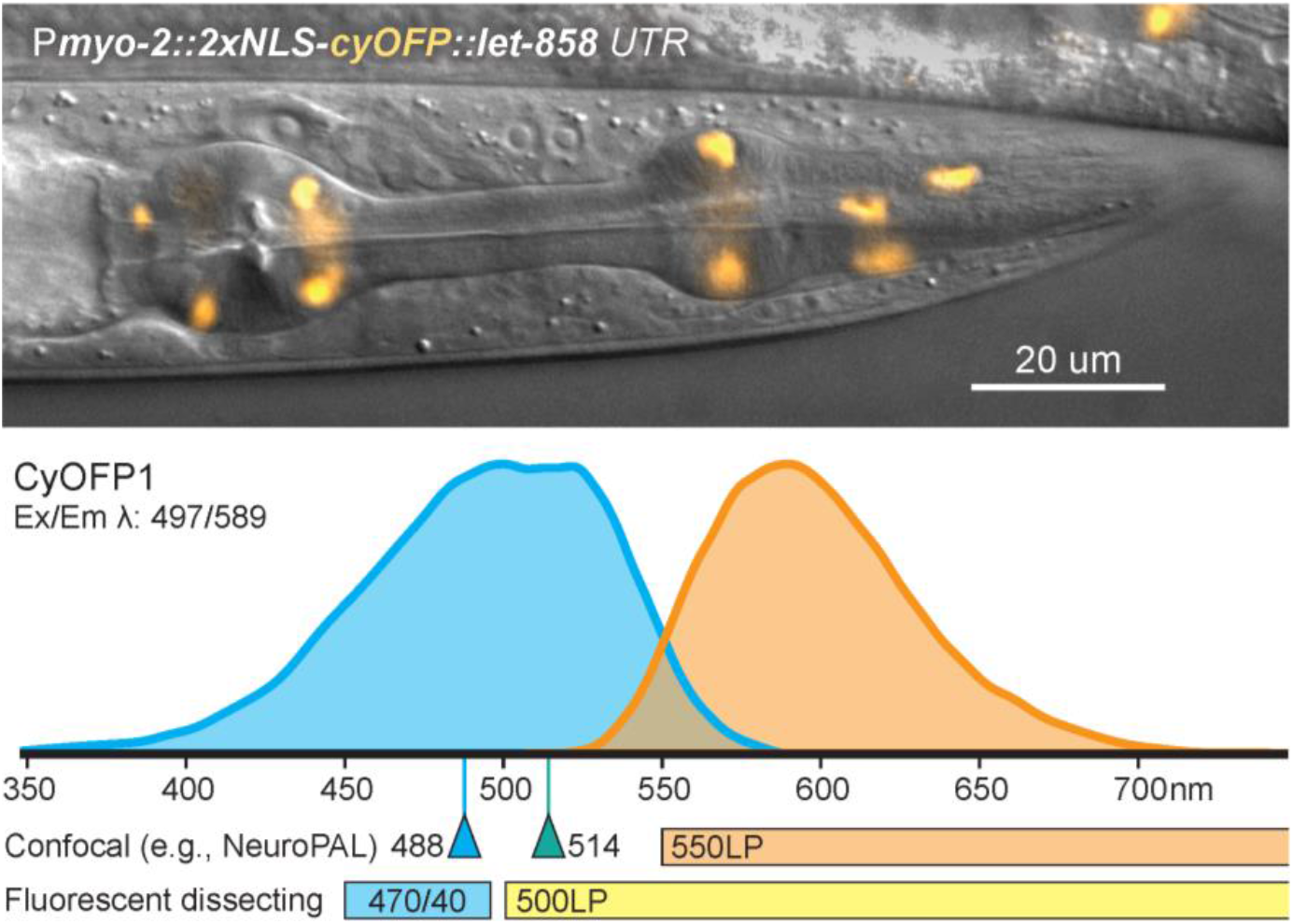
cyOFP expression pattern. (Top) Fluorescence pattern of nuclear localized ‘cyan-excitable Orange Fluorescent Protein’ in pharynx of an L4 larva from the P*myo-2::2xNLS-cyOFP::let-858 3’UTR* transgene (EG9882). Fluorescence is brightest in L3 and L4 larvae and is dim in adults. (Bottom) cyOFP has a broad excitation range (maximum from ∼480-520) and orange fluorescence emission (590 nm peak) that can be observed using a mercury lamp and standard GFP longpass filter sets on a dissecting scope or using 488 nm or 514 nm lasers with an orange-red bandpass or longpass emission filter (tagRFP, mScarlet, mRuby) on a confocal microscope. Emission and excitation spectra from fpbase.org.

**Supplemental Figure 3.**
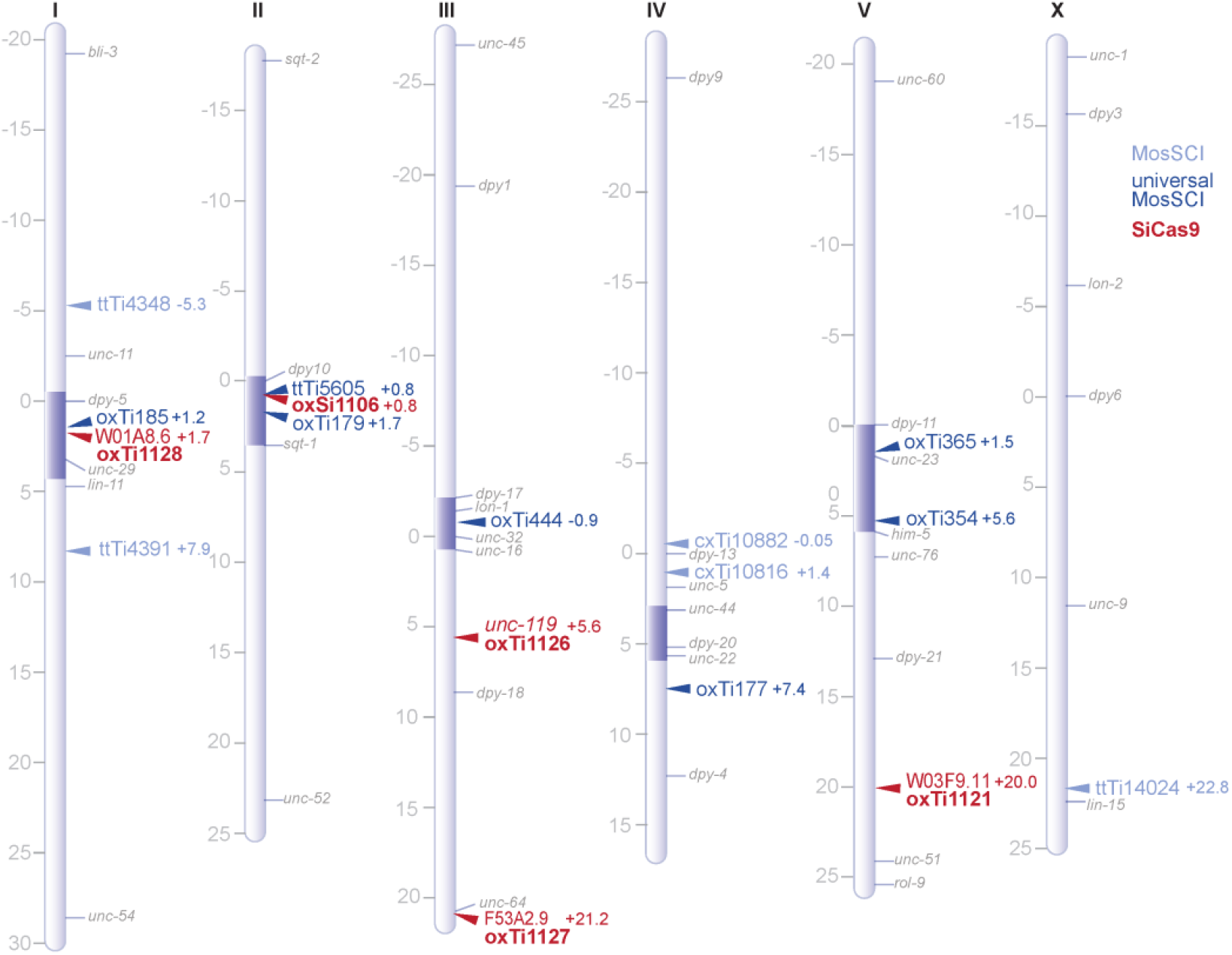
Genetic Map. Insertion sites of the integrated Cas9 alleles (red) in the genome. The locus into which the miniMos-Cas9 transposon inserted is written above (for example ‘W01A8.6’), and the original, unmodified, transposon insertions are named (‘*oxTi*’). *oxSi1106* is a single-copy insertion at the Mos1 insertion site *ttTi5605*. Also depicted are prominent MosSCI sites (light blue) and ‘universal MosSCI ‘sites (dark blue) (Frøkjær-Jensen et al., 2014, 2012). ‘Universal MosSCI’ sites use the ttTi5605 arms for homologous recombination insertion and are usually flanked by NeoR and *unc-18(+)* transgenes (see www.wormbuilder.org website).

**Supplemental Figure 4.**
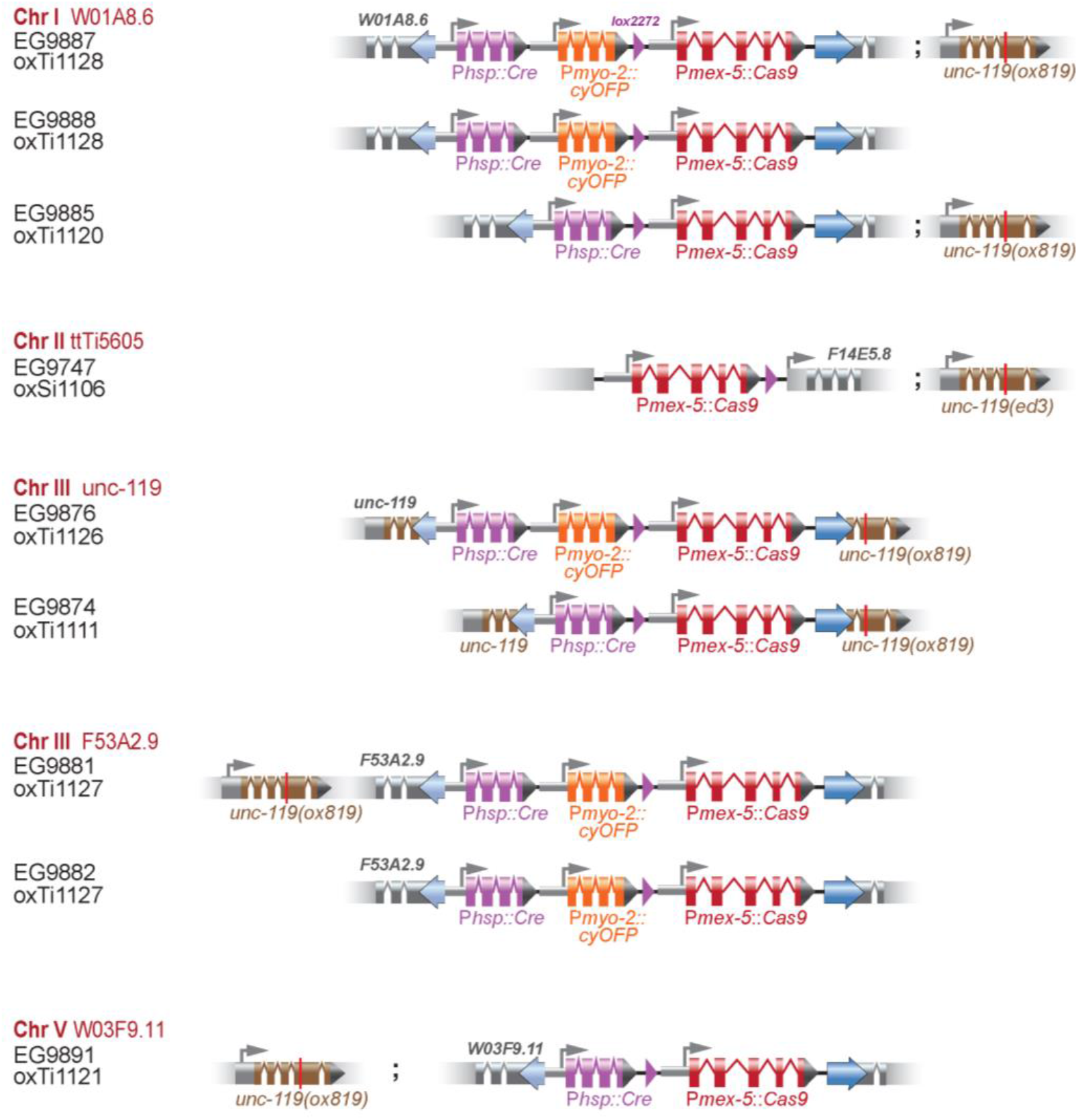
Gene models for Cas9 loci. Insertion sites are ordered by chromosome number (‘Chr’) and site into which the miniMos inserted (red). Strain and name of modified miniMos element are provided below (black). The chromosome II site is a MosSci insertion into the ttTi5605 site and does not have a Cre or orange fluorescent protein (‘cyOFP’) transgene.

**Supplemental Figure 5.**
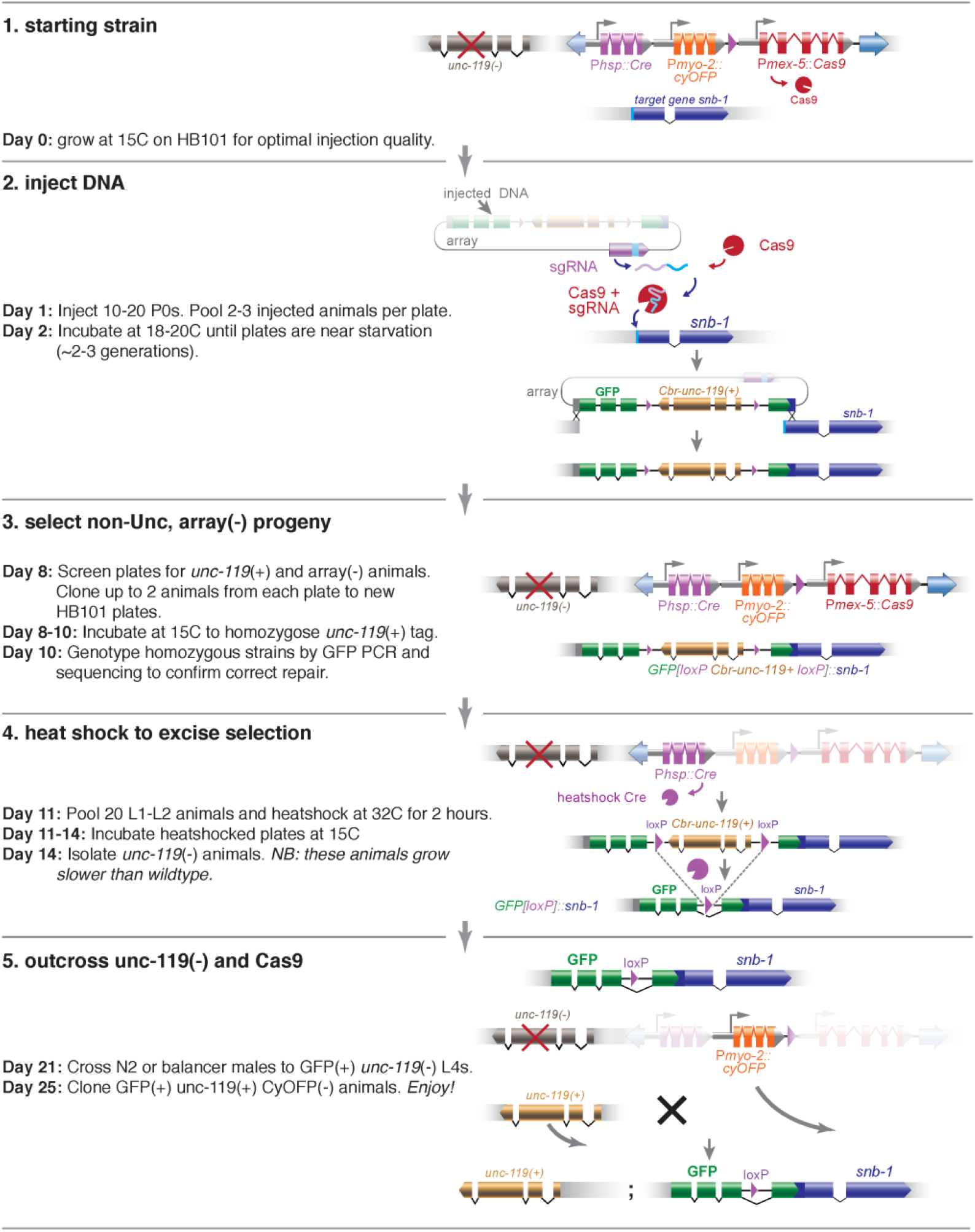
Gene tagging with GFP using integrated Cas9. Protocol for tagging *snb-1* using integrated Cas9. (1) Select a strain with integrated Cas9 that is not tightly linked to the gene otef inrest, and contains *unc-119(ox819)* if a selectable marker is desirable. (2) Inject P0s with plasmid DNAs that encode the guide RNA, your DNA template for gene tagging, and ‘red’ fluorescent markers to mark the array. (3) When P0 plates are starved, single non-Unc, non-‘Red’ animals, and confirm the presence of the edit by fluorescence or PCR. (4) Heat shock the line express Cre and excise the *Cbr-unc-119(+)* selectable marker. In the next generation, single Unc progeny. (5) Cross the strain to wild type, single heterozygotes, and then single non-orange individuals from the progeny to establish the tagged strain.

## Supplemental Tables

**Supplemental Table 1.** Post-silencing edits. Number of injected P0 worms (EG9747) that yielded at least one GFP insertion event on primary or secondary plates. ‘Primary + Secondary’ tallies P0s for which GFP+ worms were found on both the primary plate founded by the single injected P0 and on the secondary plate founded by 20 array+ GFP-progeny of the originally injected P0 (∼F3). ‘Primary Only’ tallies P0s for which GFP+ worms were found on the primary plate but not on the secondary plate. ‘Secondary Only’ tallies P0s for which GFP+ worms were not found on the primary plate but were found on the secondary plate. ‘Neither’ tallies P0s for which GFP+ worms were not found among the progeny on either the primary of secondary plate.

| Gene | Primary +<br>Secondary | Primary<br>Only | Secondary<br>Only | Neither | Total |
| --- | --- | --- | --- | --- | --- |
| *sng-1* | 1 | 4 | 1 | 2 | 8 |
| *snb-1* | 5 | 0 | 0 | 0 | 5 |
| *rab-3* | 6 | 3 | 1 | 5 | 15 |
| *unc-32* | 4 | 3 | 0 | 1 | 8 |
| *unc-17* | 19 | 0 | 0 | 0 | 19 |
| *snt-1* (short) | 0 | 0 | 0 | 6 | 6 |
| *snt-1* (long) | 4 | 2 | 2 | 13 | 21 |

**Supplemental Table 2.**
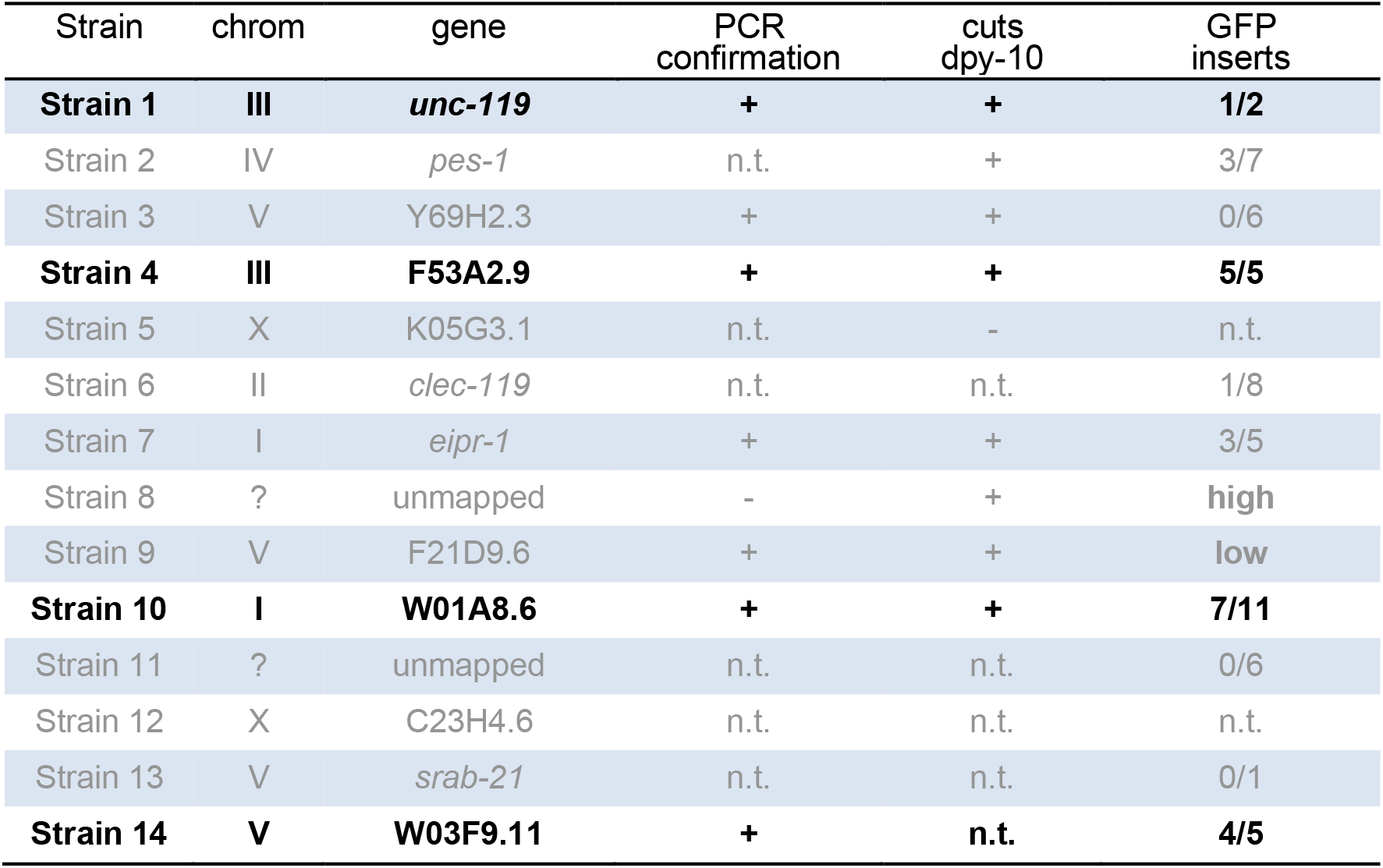
miniMos insertions. Summary of analyses of 14 Cas9 miniMos integration strains. ‘Strain’ is the isolation name. ‘chrom’ and ‘gene’ identify the location of the insert determined by inverse PCR, ‘unmapped’ means the inverse PCR failed or was ambiguous. PCR confirmation: ‘+’ indicates position was confirmed by locus-specific PCR, ‘-’ indicates confirmatory PCR failed. ‘n.t.’, not tested. ‘cuts *dpy-10*’: P0s injected with a *dpy-10* sgRNA expressing plasmid, ‘+’ indicates the presence, and ‘-’ indicates the absence of Dpy and Rol worms among the progeny of those P0s. ‘GFP inserts’: number of lines yielding *unc-32*::GFP insertions over total injected lines.

**Supplemental Table 3.** Strains. Bolded strains are the most useful for CRISPR modifications using *unc-119* selections: EG9747 chromosome II, EG9881 chromosome III, EG9887 chromosome I, EG9891 chromosome V.

| Strain | Genotype |
| --- | --- |
| N2 | - |
| EG6249 | *ttTi5605* II ; *unc-119(ed3)* III |
| none | *oxSi1196* [*Pmex-5::Cas9(cDNA)*: *tbb-2* 3'UTR *Cbr-unc-119+*; * *ttTi5605*] II ; *unc-119(ed3)* III |
| EG9615 | *oxSi1091* [*Pmex-5::cas9(+smu-2 introns)*: *tbb-2* 3'UTR *unc-119+*; * *ttTi5605*] II ; *unc-119(ed3)* III |
| **EG9747** | *oxSi1106* [*Pmex-5::Cas9:tbb-2* 3'UTR + *lox2272*; * *ttTi5605*] II ; *unc-119(ed3)* III |
| EG9814 | *unc-119(ox819)* III |
| EG9809 | *unc-119(ox819 oxTi1083* [*Pmex-5::cas9(+smu-2 introns)* + *lox2272*]) III |
| EG9873 | *unc-119(ox819 oxTi1113* [*Pmex-5::cas9(+ smu-2 introns)*, *Phsp-16.41::Cre* + *lox2272 Cbr-unc-119 lox2272*]) III |
| EG9874 | *unc-119(ox819 oxTi1111* [*Pmex-5::cas9(+ smu-2 introns)*, *Phsp-16.41::Cre* + *lox2272*]) III |
| EG9875 | *unc-119(ox819 oxTi1122* [*Pmex-5::cas9(+ smu-2 introns)*, *Phsp-16.41::Cre*, *Pmyo-2::2xNLS-CyOFP* + *lox2272 Cbr-unc-119 lox2272*]) III |
| EG9876 | *unc-119(ox819 oxTi1126* [*Pmex-5::cas9(+ smu-2 introns)*, *Phsp-16.41::Cre*, *Pmyo-2::2xNLS-CyOFP* + *lox2272*]) III |
| EG9877 | *F53A2.9(oxTi1112* [*Pmex-5::cas9(+smu-2 introns)* + *lox2272*]), *unc-119(ox819)* III |
| EG9878 | *F53A2.9(oxTi1118* [*Pmex-5::cas9(+ smu-2 introns)*, *Phsp-16.41::Cre* + *lox2272 Cbr-unc-119 lox2272*], *unc-119(ox819)* III |
| EG9879 | *F53A2.9(oxTi1118* [*Pmex-5::cas9(+ smu-2 introns)*, *Phsp-16.41::Cre* + *lox2272*]), *unc-119(ox819)* III |
| EG9880 | *F53A2.9(oxTi1123* [*Pmex-5::cas9(+ smu-2 introns)*, *Phsp-16.41::Cre*, *Pmyo-2::2xNLS-CyOFP* + *lox2272 Cbr-unc-119 lox2272*]), *unc-119(ox819)* III |
| **EG9881** | *F53A2.9(oxTi1127* [*Pmex-5::cas9(+ smu-2 introns)*, *Phsp-16.41::Cre*, *Pmyo-2::2xNLS-CyOFP* + *lox2272*]), *unc-119(ox819)* III |
| EG9882 | *F53A2.9(oxTi1127* [*Pmex-5::cas9(+ smu-2 introns)*, *Phsp-16.41::Cre*, *Pmyo-2::2xNLS-CyOFP* + *lox2272*]) III |
| EG9883 | *W01A8.6(oxTi1114* [*Pmex-5::cas9(+smu-2 introns)* + *lox2272*]) I ; *unc-119(ox819)* III |
| EG9884 | *W01A8.6(oxTi1120* [*Pmex-5::cas9(+ smu-2 introns)*, *Phsp-16.41::Cre* + *lox2272 Cbr-unc-119 lox2272*]) I ; *unc-119(ox819)* III |
| EG9885 | *W01A8.6(oxTi1120* [*Pmex-5::cas9(+ smu-2 introns)*, *Phsp-16.41::Cre* + *lox2272*]) I ; *unc-119(ox819)* III |
| EG9886 | *W01A8.6(oxTi1124* [*Pmex-5::cas9(+ smu-2 introns)*, *Phsp-16.41::Cre*, *Pmyo-2::2xNLS-CyOFP* + *lox2272 Cbr-unc-119 lox2272*]) I ; *unc-119(ox819)* III |
| **EG9887** | *W01A8.6(oxTi1128* [*Pmex-5::cas9(+ smu-2 introns)*, *Phsp-16.41::Cre*, *Pmyo-2::2xNLS-CyOFP* + *lox2272*]) I ; *unc-119(ox819)* III |
| EG9888 | *W01A8.6(oxTi1128* [*Pmex-5::cas9(+ smu-2 introns)*, *Phsp-16.41::Cre*, *Pmyo-2::2xNLS-CyOFP* + *lox2272*]) I |
| EG9889 | *unc-119(ox819)* III ; *W03F9.11(oxTi1116* [*Pmex-5::cas9(+smu-2 introns)* + *lox2272*]) V |
| EG9890 | *unc-119(ox819)* III ; *W03F9.11(oxTi1121* [*Pmex-5::cas9(+ smu-2 introns)*, *Phsp-16.41::Cre* + *lox2272 Cbr-unc-119 lox2272*]) V |
| **EG9891** | *unc-119(ox819)* III ; *W03F9.11(oxTi1121* [*Pmex-5::cas9(+ smu-2 introns)*, *Phsp-16.41::Cre* + *lox2272*]) V |
| EG9892 | *unc-119(ox819)* III ; *W03F9.11(oxTi1125* [*Pmex-5::cas9(+ smu-2 introns)*, *Phsp-16.41::Cre*, *Pmyo-2::2xNLS-CyOFP* + *lox2272 Cbr-unc-119 lox2272*]) V |
| EG9893 | *unc-119(ox819)* III ; *W03F9.11(oxTi1129* [*Pmex-5::cas9(+ smu-2 introns)*, *Phsp-16.41::Cre*, *Pmyo-2::2xNLS-CyOFP* + *lox2272*]) V |
| EG9894 | *W03F9.11(oxTi1129* [*Pmex-5::cas9(+ smu-2 introns)*, *Phsp-16.41::Cre*, *Pmyo-2::2xNLS-CyOFP* + *lox2272*]) V |

**Supplemental Table 4.** Plasmids.

| Plasmid | Description | Availability | Reference |
| --- | --- | --- | --- |
| pMLS538 | pDONR-221:2xNLS-Cas9(+*smu-2* introns) | Addgene | this study |
| pMLS539 | pDONR-221:2xNLS-Cas9(cDNA + syntron) | - | this study |
| pMLS544 | pEXPR(5605):*Pmex-5::Cas9*(+*smu-2* introns)::*tbb-2* 3'UTR; *Cbr-unc-119* | Addgene | this study |
| pMLS546 | pEXPR(5605):*Pmex-5::Cas9*(cDNA + syntron)::*tbb-2* 3'UTR; *Cbr-unc-119* | Addgene | this study |
| pMLS389 | pEXPR:*P<sub>U6</sub>::sgRNA(unc-119)**U6*-terminator | - | this study |
| pMLS597 | pEXPR:*P<sub>U6</sub>::sgRNA(dpy-10)**U6*-terminator | Addgene | this study |
| pMLS640 | pEXPR(5605):*Pmex-5::Cas9*(+*smu-2* introns)::*tbb-2* 3'UTR; lox2272 *Cbr-unc-119* lox2272 | Addgene | this study |
| pMLS621 | pDEST(5605):lox2272 *Cbr-unc-119* lox2272 | Addgene | this study |
| pMLS713 | pDEST(miniMos):lox2272 *Cbr-unc-119* lox2272 | Addgene | this study |
| pMLS714 | pEXPR(5605):*Pmex-5::Cas9*(+*smu-2* introns)::*tbb-2* 3'UTR ; lox2272 *Cbr-unc-119* lox2272 | Addgene | this study |
| pMLS715 | pDEST(miniMos-5'Arm):lox2272 *Cbr-unc-119* lox2272 | Addgene | this study |
| pMLS716 | pEXPR(miniMos-5'Arm):*Phsp16.41::Cre::tbb-2* 3'UTR ; lox2272 *Cbr-unc-119* lox2272 | Addgene | this study |
| pMLS719 | pEXPR:*P<sub>U6</sub>::sgRNA*(miniMos-5'Arm)*U6*-terminator | Addgene | this study |
| pMLS791 | pEXPR:*Phsp16.41::Cre::tbb-2* 3'UTR ; *Pmyo-2::2xNLS-cyOFP::let-858* 3'UTR ; lox2272 *Cbr-unc-119* lox2272 | Addgene | this study |
| pMLS338 | pCRISPR:*unc-32::gfp* + loxp *Cbr-unc-119* loxp | Addgene | Schwartz and Jorgensen, 2016 |
| pMLS255 | pCRISPR:*gfp::unc-17* + loxp *Cbr-unc-119* loxp | - | Schwartz and Jorgensen, 2016 |
| pSYC33 | pCRISPR:*snt-1::gfp* + loxp *Cbr-unc-119* loxp | - | this study |
| pMLS357 | pCRISPR:*sng-1::gfp* + loxp *Cbr-unc-119* loxp | - | Schwartz and Jorgensen, 2016 |
| pMLS276 | pCRISPR:*gfp::snb-1* + loxp *Cbr-unc-119* loxp | - | Schwartz and Jorgensen, 2016 |
| pMLS817 | pCRISPR:*gfp::rab-3* + loxp *Cbr-unc-119* loxp | - | this study |
| pMLS485 | pEXPR:*P<sub>eft-3</sub>::Cas9*(+*smu-2* introns)::*tbb-2* 3'UTR | Addgene | this study |

## Bibliography

Aljohani, M.D., El Mouridi, S., Priyadarshini, M., Vargas-Velazquez, A.M., Frøkjær-Jensen, C., 2020. Engineering rules that minimize germline silencing of transgenes in simple extrachromosomal arrays in C. elegans. Nat Commun 11, 6300. https://doi.org/10.1038/s41467-020-19898-0

Brenner, S., 1974. The genetics of Caenorhabditis elegans. Genetics 77, 71–94.

Chen, C., Fenk, L.A., de Bono, M., 2013. Efficient genome editing in Caenorhabditis elegans by CRISPR-targeted homologous recombination. Nucleic Acids Res 41, e193. https://doi.org/10.1093/nar/gkt805

Cho, S.W., Lee, Jihyun, Carroll, D., Kim, J.-S., Lee, Junho, 2013. Heritable gene knockout in Caenorhabditis elegans by direct injection of Cas9-sgRNA ribonucleoproteins. Genetics 195, 1177–1180. https://doi.org/10.1534/genetics.113.155853

Dickinson, D.J., Pani, A.M., Heppert, J.K., Higgins, C.D., Goldstein, B., 2015. Streamlined Genome Engineering with a Self-Excising Drug Selection Cassette. Genetics 200, 1035–1049. https://doi.org/10.1534/genetics.115.178335

Dickinson, D.J., Ward, J.D., Reiner, D.J., Goldstein, B., 2013. Engineering the Caenorhabditis elegans Genome Using Cas9-Triggered Homologous Recombination. Nat Methods 10, 1028–1034. https://doi.org/10.1038/nmeth.2641

Dokshin, G.A., Ghanta, K.S., Piscopo, K.M., Mello, C.C., 2018. Robust Genome Editing with Short Single-Stranded and Long, Partially Single-Stranded DNA Donors in Caenorhabditis elegans. Genetics 210, 781–787. https://doi.org/10.1534/genetics.118.301532

Fire, A., Alcazar, R., Tan, F., 2006. Unusual DNA Structures Associated With Germline Genetic Activity in Caenorhabditis elegans. Genetics 173, 1259–1273. https://doi.org/10.1534/genetics.106.057364

Friedland, A.E., Tzur, Y.B., Esvelt, K.M., Colaiácovo, M.P., Church, G.M., Calarco, J.A., 2013. Heritable genome editing in C. elegans via a CRISPR-Cas9 system. Nat Methods 10, 741–743. https://doi.org/10.1038/nmeth.2532

Frøkjær-Jensen, C., Davis, M.W., Ailion, M., Jorgensen, E.M., 2012. Improved Mos1-mediated transgenesis in C. elegans. Nat. Methods 9, 117–118. https://doi.org/10.1038/nmeth.1865

Frøkjaer-Jensen, C., Davis, M.W., Hopkins, C.E., Newman, B.J., Thummel, J.M., Olesen, S.-P., Grunnet, M., Jorgensen, E.M., 2008. Single-copy insertion of transgenes in Caenorhabditis elegans. Nat Genet 40, 1375–1383. https://doi.org/10.1038/ng.248

Frøkjær-Jensen, C., Davis, M.W., Sarov, M., Taylor, J., Flibotte, S., LaBella, M., Pozniakovsky, A., Moerman, D.G., Jorgensen, E.M., 2014. Random and targeted transgene insertion in Caenorhabditis elegans using a modified Mos1 transposon. Nature Methods 11, 529–534. https://doi.org/10.1038/nmeth.2889

Frøkjær-Jensen, C., Jain, N., Hansen, L., Davis, M.W., Li, Y., Zhao, D., Rebora, K., Millet, J.R.M., Liu, X., Kim, S.K., Dupuy, D., Jorgensen, E.M., Fire, A.Z., 2016. An Abundant Class of Non-coding DNA Can Prevent Stochastic Gene Silencing in the C. elegans Germline. Cell 166, 343–357. https://doi.org/10.1016/j.cell.2016.05.072

Ghanta, K.S., Mello, C.C., 2020. Melting dsDNA Donor Molecules Greatly Improves Precision Genome Editing in Caenorhabditis elegans. Genetics 216, 643–650. https://doi.org/10.1534/genetics.120.303564

Kelly, W.G., Xu, S., Montgomery, M.K., Fire, A., 1997. Distinct requirements for somatic and germline expression of a generally expressed Caernorhabditis elegans gene. Genetics 146, 227–238.

Mello, C.C., Kramer, J.M., Stinchcomb, D., Ambros, V., 1991. Efficient gene transfer in C.elegans: extrachromosomal maintenance and integration of transforming sequences. EMBO J 10, 3959–3970.

Norris, A.D., Kim, H.-M., Colaiácovo, M.P., Calarco, J.A., 2015. Efficient Genome Editing in Caenorhabditis elegans with a Toolkit of Dual-Marker Selection Cassettes. Genetics 201, 449–458. https://doi.org/10.1534/genetics.115.180679

Paix, A., Folkmann, A., Rasoloson, D., Seydoux, G., 2015. High Efficiency, Homology-Directed Genome Editing in Caenorhabditis elegans Using CRISPR-Cas9 Ribonucleoprotein Complexes. Genetics 201, 47–54. https://doi.org/10.1534/genetics.115.179382

Schaner, C.E., Kelly, W.G., 2006. Germline chromatin. WormBook. https://doi.org/10.1895/wormbook.1.73.1

Schwartz, M.L., Jorgensen, E.M., 2016. SapTrap, a Toolkit for High-Throughput CRISPR/Cas9 Gene Modification in Caenorhabditis elegans. Genetics 202, 1277–1288. https://doi.org/10.1534/genetics.115.184275

Stinchcomb, D.T., Shaw, J.E., Carr, S.H., Hirsh, D., 1985. Extrachromosomal DNA transformation of Caenorhabditis elegans. Molecular and Cellular Biology 5, 3484–3496. https://doi.org/10.1128/MCB.5.12.3484

Waaijers, S., Portegijs, V., Kerver, J., Lemmens, B.B.L.G., Tijsterman, M., van den Heuvel, S., Boxem, M., 2013. CRISPR/Cas9-Targeted Mutagenesis in Caenorhabditis elegans. Genetics 195, 1187–1191. https://doi.org/10.1534/genetics.113.156299

Wang, H., La Russa, M., Qi, L.S., 2016. CRISPR/Cas9 in Genome Editing and Beyond. Annu. Rev. Biochem. 85, 227–264. https://doi.org/10.1146/annurev-biochem-060815-014607

Zhang, D., Tu, S., Stubna, M., Wu, W.-S., Huang, W.-C., Weng, Z., Lee, H.-C., 2018. The piRNA targeting rules and the resistance to piRNA silencing in endogenous genes. Science 359, 587–592. https://doi.org/10.1126/science.aao2840

